# Statistically-estimated tree biomass, stem density, and basal area for the upper Midwestern United States at the time of Euro-American settlement

**DOI:** 10.1101/856526

**Authors:** Christopher J. Paciorek, Charles V. Cogbill, Jody A. Peters, Simon J. Goring, John W. Williams, David J. Mladenoff, Andria Dawson, Jason S. McLachlan

## Abstract

We present gridded 8 km-resolution data products of the estimated biomass, basal area, and stem density of tree taxa at the time of Euro-American settlement of the midwestern United States for the states of Minnesota, Wisconsin, Michigan, Illinois, and Indiana. The data come from settlement-era Public Land Survey (PLS) data (ca. 0.8-km resolution) of trees recorded by land surveyors. The surveyor notes have been transcribed, cleaned, and processed to estimate biomass, basal area, and stem density at individual points on the landscape. The point-level data are then aggregated within grid cells and statistically smoothed using a statistical model that accounts for zero-inflated continuous data with smoothing based on generalized additive modeling techniques and approximate Bayesian uncertainty estimates. We expect this data product to be useful for understanding the state of vegetation in the midwestern United States prior to large-scale Euro-American settlement. In addition to specific regional questions, the data product can serve as a baseline against which to investigate how forests and ecosystems change after intensive settlement. The data products (including both raw and statistically smoothed estimates at the 8-km scale) are being made available at the LTER network data portal as version 1.0.

## Introduction

Terrestrial vegetation in midwestern North America changed drastically at the time of Euro-American settlement (McAndrews 1988; Rhemtulla et al. 2009). Before settlement, the midwestern United States was the location of a major ecological transition between the grasslands of the Great Plains and the forests of eastern and northern North America (Transeau 1935, Grimm 1984, Danz et al. 2013). These grasslands have now mostly been replaced by agriculture or pastoral land use, except in areas of prairie conservation or restoration. Forested areas were also heavily affected by clearance for agriculture and logging during and after settlement (Rhemtulla et al. 2009; Schulte et al. 2007, Goring et al. 2016). Historical datasets from this time period, collected during the time of land surveys and allotment, provide critical context for understanding terrestrial ecology, the carbon cycle, and vegetation-atmosphere feedbacks (Caspersen et al. 2000, Rhemtulla et al. 2009, Lawrence et al. 2016). They allow researchers to define ‘baseline’ conditions for purposes of conservation planning, to understand ecosystem processes at decadal and centennial scales, to track how vegetation changes with changing climate, and to understand changes in ecosystems after widespread land use change. Here we present methods for and statistical estimates of biomass, basal area, and stem density on an 8 km grid across a set of states in the midwestern United States.

Euro-American settlement and subsequent land use change occurred over many decades across North America. During that time, land surveys were done to demarcate land for land tenure and use, usually involving recording and marking trees adjacent to survey corners. These data provide vegetation information that can be mapped and used quantitatively to represent forest composition, and, sometimes, structure at the period of settlement. In the northeastern United States, early surveys provide only data at the township level (Cogbill et al. 2002, Thompson et al. 2013), which cannot be used to estimate biomass, basal area, or stem density, but which we have used to estimate composition (Paciorek et al. 2016). Later surveys after the establishment of the U.S. Public Land Survey System (PLS) by the General Land Office (GLO) provide point-level (i.e., corner-level) data along a regular grid, every one-half mile (800 m) spacing, for Ohio and westward during the period 1785 to 1907 (Bourdo 1956, Pattison 1957, Schulte and Mladenoff 2001, Goring et al. 2016). At each point 2-4 trees were identified, and the common name, diameter at breast height (dbh), and distance and bearing from the point to the trees were recorded. Using statistical techniques (Cogbill et al. 2018), these data allow us to estimate biomass, basal area, and stem density at each point. These point-level data are quite noisy, but can be aggregated to coarser spatial resolution to more robustly estimate spatial patterns of vegetation (Goring et al. 2016). At the 8 km grid resolution, the estimates are still noisy, and there are some spatial gaps in the available data, so in this work we employ a spatial statistical model to smooth over the noise and impute in missing grid cells. The result is a statistical data product that provides statistical estimates of biomass and density with quantitative estimates of uncertainty.

In contrast to Paciorek et al. (2016) we estimate biomass and density rather than composition and we use an extended dataset with additional data cleaning steps that were applied more consistently across the region. Relative to Goring et al. (2016) we use a spatial statistical model to smooth over the noisy grid-level estimates; we extend the domain to include southern Michigan, Illinois, and Indiana; we use updated allometric scaling factors from Chojnacky et al. (2014); and we apply additional and more consistent data cleaning steps across the domain.

In Section Methods, we describe the procedures used to obtain and clean the PLS survey data at the survey points, followed by processing to homogenize the data across the states of interest, and finally the statistical methodology used to estimate biomass, basal area, and stem density, first at the individual survey points, and then on an 8 km by 8 km grid, stratified by taxon and in total. In Section Results we present the results of cross-validation work carried out to determine the optimal statistical smoothing approach, and present basic summaries of biomass and stem density. In Section Data Product we describe the various data products we have produced and archived. Finally, in Section Discussion we discuss the uncertainties estimated by the statistical model and the limitations of the model.

## Methods

### PLS data collection and cleaning

The PLS was developed to enable the division and sale of federal lands from Ohio westward. The survey created a 1 mile^2^ (2.56 km^2^) grid (sections) on the landscape. At each half-mile (quarter-section) and mile (section) survey point, a post was set or a tree was blazed as the official location marker. PLS surveyors then recorded tree stem diameters, measured distances and bearings of the two to four trees adjacent to the survey point, and identified tree taxa using common (and often regionally idiosyncratic) names. In the Midwest, PLS data thus represent measurements by hundreds of surveyors from 1786 until 1907, with changing sets of instructions over time (Stewart 1935, White 1983). Survey procedures varied widely in Ohio and distance, diameter, and bearing information are not systematically available, so Ohio is not included in this work. The work presented here builds upon prior digitization and classification of PLS data for Wisconsin, Minnesota, and Michigan, with extensive additional cleaning and correction of the Michigan data and extensive additional digitization of Illinois and Indiana by the authors. Digitization of PLS data in Minnesota, Wisconsin and Michigan’s Upper Peninsula and northern Lower Peninsula is essentially complete, with PLS data for nearly all 8 km grid cells. Data for the southern portion of Michigan’s Lower Peninsula include the section points, but the quarter-section points have not been digitized yet except for the Detroit region, which is complete. Data in Illinois and Indiana represent a sample of the full set of grid cells, with survey record transcription ongoing at Notre Dame (see Fig. 1 for data availability).

**Fig. 1:**
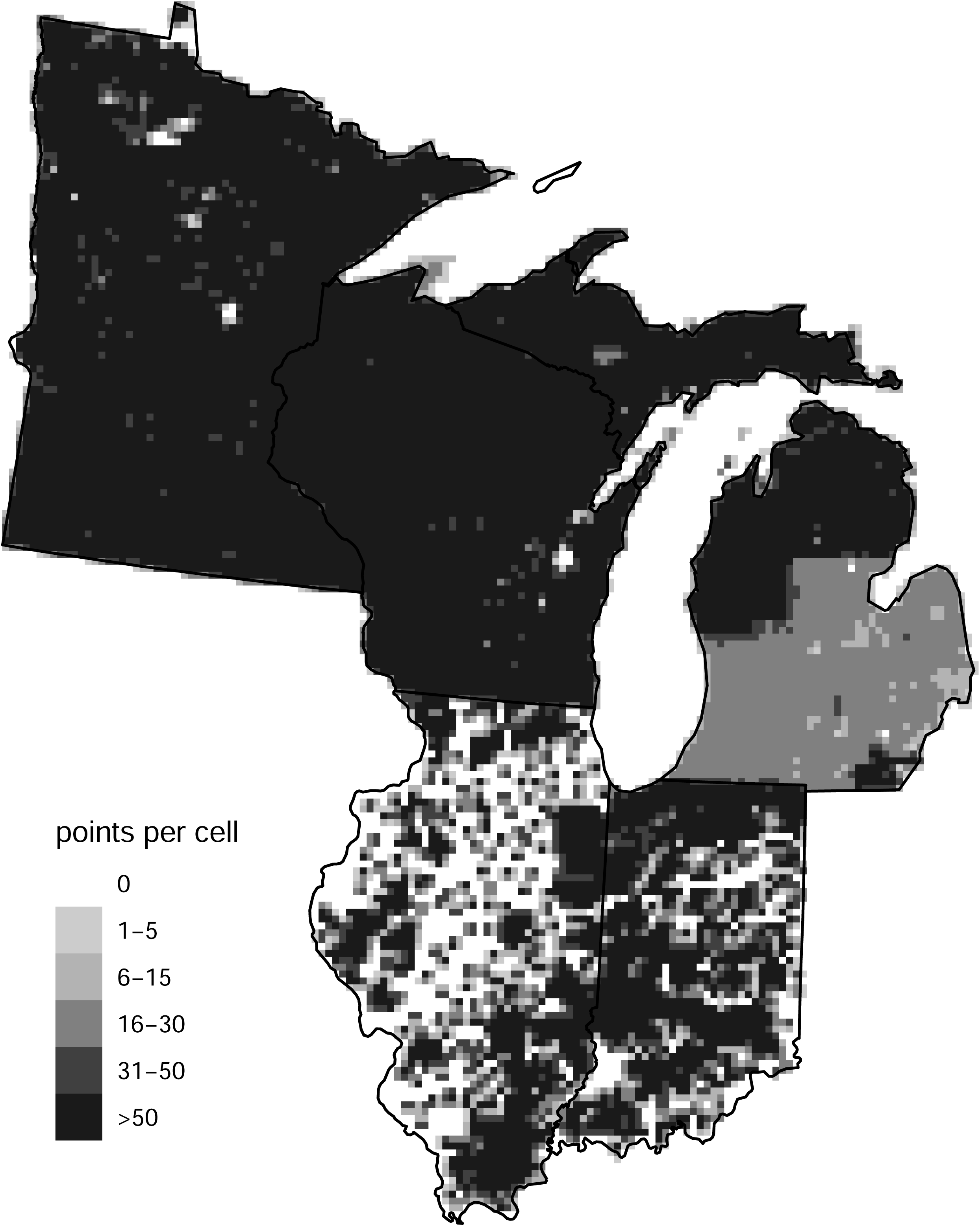
Number of PLS points per 8-km grid cell used for statistical estimation. Lighter grey in southern Michigan is caused by lack of quarter-section points. Illinois and Indiana digitization is ongoing.

As discussed in Paciorek et al. (2016), the surveys in our domain occurred over a period of more than 100 years (starting in 1799 in Indiana and ending in 1907 in Minnesota) as settlers from the United States and Europe settled what is now the midwestern United States. Our estimates are for the period of settlement represented by the survey data and therefore are time-transgressive; they do not represent any single point in time across the domain, but rather the state of the landscape at the time just prior to widespread Euro-American settlement and land use (Whitney, 1996; Cogbill et al., 2002). These datasets do include the effects of Native American land use and early Euro-American settlement activities (e.g., Black et al., 2006), but it is likely that the imprint of this earlier land use is locally concentrated rather than spatially extensive (Munoz et al., 2014).

We used expert judgment (co-author CVC) and prior work to determine the current common names of surveyor-recorded vernacular terms and abbreviations of settlement-era common names. We then aggregated into taxonomic groups that are primarily at the genus level but include some monospecific genera. We use the following 20 taxa plus an “other hardwood” category: Ash (*Fraxinus spp.*), Basswood (*Tilia americana*), Beech (*Fagus grandifolia*), Birch (*Betula spp.*), Black gum/sweet gum (*Nyssa sylvatica* and *Liquidambar styraciflua*), Cedar/juniper (J*uniperus virginiana* and *Thuja occidentalis*), Cherry (*Prunus spp.*), Dogwood (*Cornus spp.*), Elm (*Ulmus spp.*), Fir (*Abies balsamea*), Hemlock (*Tsuga canadensis*), Hickory (*Carya spp.*), Ironwood (*Carpinus caroliniana* and *Ostrya virginiana*), Maple (*Acer spp.*), Oak (*Quercus spp.*), Pine (*Pinus spp.*), Poplar/tulip poplar (*Populus spp.* and *Liriodendron tulipifera*), Spruce (*Picea spp.*), Tamarack (*Larix laricina*), and Walnut (*Juglans spp.*). Note that because of several cases of ambiguity in the common tree names used by surveyors (black gum/sweet gum,ironwood, poplar/tulip poplar, cedar/juniper), a group can represent trees from different genera or even families.

In Appendix A, we describe the specific data cleaning steps we applied to each sub-dataset as well as a variety of steps to standardize the dataset across states and minimize the potential effects of surveyor bias upon estimates of vegetation. Note that the division between northern and southern Michigan is caused by obtaining the data from different sources and can be seen in the black to grey transition in Fig. 1. In the remainder of this section, we briefly describe some of these key cleaning and standardization steps.

Following Goring et al. (2016), we excluded line and meander trees (i.e., trees encountered along survey lines as compared to trees located at section or quarter-section corners). Surveyor selection biases for tree size and species appear to have been more strongly expressed for line trees. Meander trees were used to avoid obstacles, such as water-bodies, and so have non-random habitat preferences (Liu et al. 2011).

We attempted to exclude points in water including points with information indicating wetlands without trees present; the specifics of how we did this varied by state as described in detail in Appendix A. Relative to Goring et al. (2016) we excluded some additional points in Wisconsin and Minnesota based on information indicating the presence of standing water. Note that points with a single tree might be in areas with low tree density such that the second tree was too far for the surveyors to mark it. However, in some cases a single tree may be marked because three of the quadrants were inaccessible, generally due to wet conditions. We generally excluded one-tree points based on information suggesting this was a result of water rather than because of low tree density, but in many cases it was hard to distinguish between these cases.

Relative to Goring et al. (2016) we carried out extensive additional quality control of the northern Michigan data, based on which we detected and subsequently fixed anomalies in the data. We also excluded a number of additional points with no tree data where the data were judged to be unreliable rather than indicative of low density.

As part of the overall PalEON project, we have been digitizing the PLS data from southern Michigan, Illinois, and Indiana. Goring et al. (2016) did not analyze southern Michigan, Illinois, or Indiana, while Paciorek et al. (2016) did analyze these areas, but used an earlier version of the PLS dataset only to estimate composition. The southern Michigan data were digitized from Mylar maps and found to have a variety of errors. For this work, based on the original field notes for southern Michigan, we fixed errors in some points and excluded some points with data judged to be unreliable. For Indiana and Illinois, we now have additional digitized PLS data that was unavailable in Paciorek et al. (2016)

### Estimation of point-level density

We estimated stem density at each point with a Morisita plotless density estimator that uses the measured distances from each survey point to the nearest trees at the point location (Cogbill et al. 2018). The standardized approach for the Morisita method is well-validated. However, over time the survey design used by PLS surveyors changed as protocols were updated, which affects how we estimate density from the information at each point. Appendix B summarizes the changes in the information recorded and how we developed and applied correction factors to the Morisita estimator (Goring et al. 2016, Cogbill et al. 2018) to account for these changes when estimating stem density at a point.

We limited estimates of density to trees above 8 inches dbh because that is approximately the size below which surveyors tended to avoid sampling small trees. However, in many cases smaller trees were reported by surveyors. We included all trees that were surveyed in our initial density estimate (including those with missing diameters), giving a raw stem density estimate whose meaning (in terms of the implicit diameter threshold) varies spatially based on how surveyors selected for tree size in a given area. We then used a correction factor (see Goring et al. (2016) and Cogbill et al., in progress) to scale the raw density estimates based spatially-varying estimates of the diameter distributions of PLS trees. This gives us a corrected stem density estimate for trees greater than 8 inches dbh.

Distances from the tree to the survey point were taken to be the distance from the survey notes plus one-half the diameter of the tree.

We used bearing angle information to screen and correct for points where surveyors may not have followed the PLS instructions. Specifically, we searched for and found 9602 four-tree points where the two nearest trees fell in the same quadrant. We excluded these points as this indicates the surveyors did not follow the survey instructions, and there is no rigorous way to use the Morisita estimator to estimate density for these points. In cases where information on the quadrant was missing for one or both trees we assumed they fell in different quadrants and did calculate stem density.

We removed 3629 points with one tree at a distance of zero or missing distance as it is unclear what density to estimate for such points. Many of these corners have a single corner tree with a zero distance (presumably a “corner tree” used as the corner post), which our density estimator would assign a density of infinity. We removed 1821 points with two trees and either distance missing. We also removed 131 points with two trees at distance zero. Note that points with one of two trees at distance zero do allow estimation of density using the Morisita estimator and were included.

We estimated density for one-tree points as 0.3146 stems per hectare. This density estimate is equal to one tree in a circle of radius equal to 500 links (approximately 100 m), which approximates how far a surveyor might have gone to find a second tree. Surveyors were instructed to find two trees, so the presence of only one tree generally indicates low density. While 500 links is arbitrary, our results should be insensitive to the exact value of the near-zero density that we use in such cases.

We truncated estimated densities at 10,000 stems per hectare (one tree per square meter) to reduce the influence of outlying high density values, truncating 139 points when estimating stem density itself and 246 points when estimating stem density for the biomass (and basal area) estimation (i.e., omitting the scaling to trees greater than 8 inches dbh as discussed below).

After all removals we estimated stem density at 66,648 Illinois points, 67,072 Indiana points, 113,801 Michigan points, 226,047 Minnesota points, and 159,058 Wisconsin points (Fig. 1).

### Estimation of point-level biomass

#### Estimation of individual tree biomass

We use the aboveground biomass (AGB, component 2, dry, live stump, stem, branches and foliage relationships provided in Chojnacky et al. (2014)) to estimate aboveground biomass. The assignment of allometric coefficients (for simple linear regressions of log biomass (kg) on log dbh (cm)) to taxa is provided in our Github repository (https://github.com/PalEON-Project/PLS_products) and in Table 1. Note that some of the 21 taxa use the same allometric equations.

**Table 1:** Allometric scaling coefficients used to estimate (log) biomass (kg) from (log) dbh (cm).

| PLS taxonomic grouping | Chojnacky et al. (2014) group | intercept | slope |
| --- | --- | --- | --- |
| Alder | Betulaceae < 0.40 spg | -2.5932 | 2.5349 |
| Ash | Oleaceae ≥ 0.55 spg | -1.8384 | 2.3524 |
| Atlantic white cedar | Cupressaceae < 0.30 spg | -1.9615 | 2.1063 |
| Bald cypress | Cupressaceae ≥ 0.40 spg | -2.6327 | 2.4757 |
| Basswood | Hippocastanaceae/Tiliaceae | -2.4108 | 2.4177 |
| Beech | Fagaceae, deciduous | -2.0705 | 2.441 |
| Birch | Betulaceae 0.50–0.59 spg | -1.8096 | 2.348 |
| Black gum | Cornaceae/Ericaceae/Lauraceae/Platanaceae/Rosaceae/Ulmaceae | -2.2118 | 2.4133 |
| Black gum/sweet gum | Cornaceae/Ericaceae/Lauraceae/Platanaceae/Rosaceae/Ulmaceae | -2.2118 | 2.4133 |
| Buckeye | Hippocastanaceae/Tiliaceae | -2.4108 | 2.4177 |
| Cedar/juniper | Cupressaceae < 0.30 spg | -1.9615 | 2.1063 |
| Cherry | Cornaceae/Ericaceae/Lauraceae/Platanaceae/Rosaceae/Ulmaceae | -2.2118 | 2.4133 |
| Chestnut | Fagaceae, deciduous | -2.0705 | 2.441 |
| Dogwood | Cornaceae/Ericaceae/Lauraceae/Platanaceae/Rosaceae/Ulmaceae | -2.2118 | 2.4133 |
| Elm | Cornaceae/Ericaceae/Lauraceae/Platanaceae/Rosaceae/Ulmaceae | -2.2118 | 2.4133 |
| Fir | Abies < 0.35 spg | -2.3123 | 2.3482 |
| Hackberry | Cornaceae/Ericaceae/Lauraceae/Platanaceae/Rosaceae/Ulmaceae | -2.2118 | 2.4133 |
| Hemlock | Tsuga < 0.40 spg | -2.348 | 2.3876 |
| Hickory | Fabaceae/Juglandaceae, Carya | -2.5095 | 2.6175 |
| Ironwood | Betulaceae ≥ 0.60 spg | -2.2652 | 2.5349 |
| Locust | Fabaceae/Juglandaceae, other | -2.5095 | 2.5437 |
| Maple | Aceraceae ≥ 0.50 spg | -1.8011 | 2.3852 |
| Mulberry | Fagaceae, deciduous | -2.0705 | 2.441 |
| Oak | Fagaceae, deciduous | -2.0705 | 2.441 |
| Other hardwood | average of all hardwoods | -2.2424 | 2.4498 |
| Pine | Pinus < 0.45 spg | -2.6177 | 2.4638 |
| Poplar | Salicaceae < 0.35 spg | -2.6863 | 2.4561 |
| Poplar/tulip poplar | Magnoliaceae | -2.5497 | 2.5011 |
| Spruce | Picea ≥ 0.35 spg | -2.1364 | 2.3233 |
| Sweet gum | Cornaceae/Ericaceae/Lauraceae/Platanaceae/Rosaceae/Ulmaceae | -2.2118 | 2.4133 |
| Sycamore | Cornaceae/Ericaceae/Lauraceae/Platanaceae/Rosaceae/Ulmaceae | -2.2118 | 2.4133 |
| Tamarack | Larix | -2.3012 | 2.3853 |
| Tulip poplar | Magnoliaceae | -2.5497 | 2.5011 |
| Walnut | Fabaceae/Juglandaceae, Carya | -2.5095 | 2.6175 |
| Willow | Salicaceae ≥ 0.35 spg | -2.4441 | 2.4561 |

Our original goal was to make use of the full set of allometric information in Jenkins et al. (2003) and Chojnacky et al. (2014) to incorporate uncertainty in scaling dbh to tree biomass, using the Bayesian statistical methods provided in the PEcAn software (Dietze et al. 2013) allometry module. However, even at the taxonomic aggregation inherent in our 21 taxa, there are often few allometries available for a given one of the taxonomic groupings and in many cases the allometries come from locations outside of our midwestern US spatial domain. Furthermore, although there are more allometries for stem biomass (component 6; note that this excludes branches) than for aboveground (component 2) or total biomass (component 1), most research focuses on aboveground biomass rather than stem biomass. As a result we felt that we could not robustly estimate the aboveground biomass allometries with uncertainty and have omitted this.

#### Estimation of point-level biomass and basal area

Here we describe how we biomass at each PLS point. Calculations for basal area are equivalent.

In the usual case of having two trees, we calculated the point-level biomass using one-half the stem density multiplied by the estimated biomass of each tree. When the two trees were of different taxa, this produces point-level biomass for two taxa that were added to estimate total biomass. When of the same taxa, this is equivalent to averaging the tree-level biomass for the two trees and multiplying by stem density.

For simplicity we excluded all 3221 points with any tree-level missing biomass values (i.e., missing diameters), although we note that it is possible to estimate (1) total biomass based on having one of two trees with available biomass and (2) taxon-level biomass from the available tree. The exclusion puts two-tree points on a similar footing with one-tree points (for which missing biomass prevents estimation) with the goal of limiting bias at the grid cell level.

When estimating biomass, we used the original density estimate without using correction factors that scale to the density of trees greater than 8 inches dbh. This is necessary since the trees’ biomass can only be calculated based on theoriginal density. Thus the original density combined with biomass estimates for all individual trees (including those less than 8 inches dbh) gives an unbiased biomass estimate without an explicit size threshold. We recognize this introduces some imprecision, but we note that given the limited contribution of smaller trees to total biomass, the presence or absence of a diameter threshold should have minimal effect. In contrast, for density estimation it is critical to define a threshold in order to have a meaningful quantity.

### Statistical modeling at the grid scale

#### Grid-level estimation

Before doing the statistical modeling at the 8-km grid scale, we aggregated the point-level data to the 8-km grid by averaging over point-level biomass, basal area, and stem density values for all points in a grid cell. In addition, for our statistical modeling to best account for the high abundance of points with either no trees or (for taxon-specific analyses) no trees of a given taxon, we also calculated the proportion of points in each grid cell with no trees (for taxon-specific analysis, the proportion of points with no trees of the taxon of interest).

Note that given heterogeneity in density values within a grid cell (i.e., density varies by stand), our estimates at the grid level must account for the species-density relationship. Traditionally, basal area has been calculated as the product of the mean density and mean tree basal area, but because of their negative correlation, this overestimates the average values (Bouldin 2008, 2010; Kronenfeld 2015). Therefore, our estimates of biomass and basal area in a grid cell is the mean of the point-level multiplication of density and tree size (Cogbill et al. 2018). Similarly our estimate of density for a given taxon is equal to the average of the point-level density estimates for that taxon, not to the taxon proportion of stems in a grid cell multiplied by the grid cell estimate of total density. Furthermore, the biomass of each taxon is not estimated from the taxon proportion multiplied by the overall biomass, but the mean of the point-level biomass estimates.

#### Statistical smoothing

The major challenge of modeling biomass, basal area, and stem density data is that these quantities are both non-negative and continuous, with a discrete spike at zero; few statistical distributions are available for this type of data. The description below is specifically for biomass for concreteness and clarity of presentation, but the modeling structure is the same for basal area and stem density.

There are many zero-inflated models in the statistical literature, most focusing on count or proportional data (Lambert 1992, Hall 2000). In early efforts we considered a Tweedie model (Tweedie 1984, Jørgensen, 1987) to deal with our zero-inflated continuous data. However, computational difficulties affected model convergence and the Tweedie model resulted in poor fits. Given this we developed a two-stage model to address the challenge of zero inflation in non-negatively valued distributions. Our model was motivated by the biological insight that local conditions may prevent a taxon from occurring in an area even though the taxon may be present at high density nearby. Thus we combine a model for “potential biomass”, which reflects the large-spatial-scale patterns in biomass with a model for “occupancy”, which reflects the propensity for a given forest stand to contain the taxon. This model allows for zero inflation because a low value of the probability of occupancy can easily produce observations that are zero at the grid cell aggregation.

Let *N* (*s*) be the number of PLS sample points in grid cell *s*. Let *n*_*p*_(*s*) be the number of points in grid cell *s* that have one or more trees of taxon *p*. Let 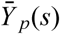 be the average biomass for taxon *p* calculated only from the *n*_*p*_(*s*) points at which the taxon is present. In other words,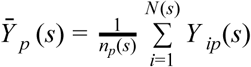 where *i* indexes the *N* (*s*) sample points within cell *s*. Let *m*_*p*_(*s*) be the potential (log) biomass process and θ_*p*_(*s*) be the occupancy process, both evaluated at grid cell *s*. The biomass in a grid cell can then be calculated as *b*_*p*_(*s*) = θ_*p*_(*s*) exp(*m*_*p*_(*s*)), namely weighting the average biomass in “occupied points” by the proportion of points that contain the taxon.

First consider the occupancy model. The likelihood is binomial, *n*_*p*_(*s*) ∼ *Bin*(*N* (*s*), θ_*p*_(*s*)). Note that the occupancy model represents the occupancy of points within a grid cell for taxon *p* and that 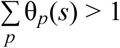 because two taxa will often “occupy” the same point, since most PLS points have two trees. Next consider the (log) biomass process. We considered modeling potential biomass both on the original scale, 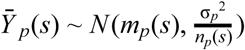, and on the log scale, 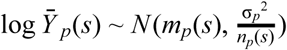, where the scaling of the variance by *n*_*p*_(*s*) is the usual variance of an average. Note that this likelihood accounts for heteroscedasticity related to the number of points at which the taxon is observed (not the number of PLS points in the cell). Finally, for total (non-taxon-specific) biomass, *n*_*p*_(*s*) above is simply the number of points with any trees.

However, based on the delta method, the correct approximate distribution when working on the log scale is 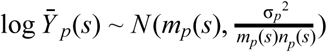. There is no clear means of accounting for the extra *m*_*p*_(*s*) in the denominator when fitting the potential biomass on the log scale using generalized additive modeling software (see below). Despite this we found that working on the log scale produced more accurate point and uncertainty estimates based on cross-validation. This improved performance likely results from i) downweighting the influence of outliers and ii) the log-scale model inherently having the variance scale with the mean (when both are considered on the original scale), which we observe empirically in the raw grid-level data.

This two-stage model is able to account for the zero inflation produced by structural zeros (the taxon is not present because local conditions prevent it) through the use of the occupancy model. Through the potential model, it is also able to capture the smooth larger-scale variation in biomass. And by having both component models, we can account for the differential amounts of information in the face of the large number of zeros and different numbers of sampling points in each grid cell.

Note that *m*_*p*_(*s*) is likely to be quite smooth spatially, at least for the PLS data, because when a point is occupied by a given taxon, the tree is likely to be of adult size, regardless of whether the tree is common in the grid cell. So most of the spatial variation in biomass may be determined by variability in occupancy. The potential biomass is meant to correct for the fact that density and tree size may vary somewhat, but probably not drastically, across the domain.

We fit the two component models using penalized splines to model the spatial variation, with the fitting done by the numerically robust generalized additive modeling (GAM) methodology implemented in the R package mgcv (Wood 2017), using the GAM implementation intended for large datasets encoded in the bam() function (Wood et al. 2015) in place of the usual gam() function.

We accounted for the heterogeneity in the number of occupied points per grid cell by setting the ‘weights’ argument in the bam() function to equal to *n*_*p*_(*s*). We also considered scaling all weights by dividing by 70, where 70 is the approximate number of points in a cell that was fully-surveyed. This treats a fully-covered cell as having one ‘unit’ of information and scales the contribution to the likelihood from cells with a different number of points relative to that. However, the results with and without the division by 70 were identical for the point estimates and very similar for the uncertainty estimates, so our final results omit this scaling.

We did not use covariates as predictors in our statistical model for several reasons. First we have fairly complete coverage (see Fig. 1), such that the use of covariates is expected to provide limited additional information. Second, covariates such as climate for the settlement time period are not available and we were reluctant to make assumptions that present-day values are sufficiently similar to values in the past. Finally, without developing complicated statistical models that allow the effect of covariates to vary spatially (so-called varying coefficient models), using regression coefficient estimates that are constant spatially can cause biases, such as inferring the presence of a taxon outside of its range boundary. For these reasons, we chose to rely only on spatial smoothing of the raw data. Future researchers could use our raw data products in combination with covariates.

Finally, to estimate total biomass, we fit the model above to raw total biomass values from the survey points, aggregating in the same fashion as described above for individual taxa, but including data from all trees.

#### Quasi-Bayesian uncertainty estimates

As discussed in Wood (2017), one can derive a quasi-Bayesian approach and simulate draws from an approximate Bayesian posterior by drawing values of the spline coefficients based on the approximate Bayesian posterior covariance provided by gam() or bam() and, for each draw, calculating a draw of θ_*p*_(*s*) and similarly a draw for *m*_*p*_(*s*) for the biomass process. We combined 250 draws from the occupancy and potential biomass processes (assuming independence between the processes) to produce biomass draws for each taxon and for total biomass. The procedure for stem density was analogous.

Note that one major drawback of this methodology is that individual taxon estimates are not constrained to add to the total biomass values estimated from using our model on raw total biomass values because the taxa are fit individually. Further, as was the case in our related modeling of composition (Paciorek et al. 2016), we do not capture correlations between taxa, in part to reduce computational bottlenecks and in part to avoid inferring the value of one taxon based on the value of another. While there are real correlations, the correlation structure likely varies substantially over space (e.g., two taxa that covary strongly can have different range boundaries such that the presence of one beyond the boundary of another does not indicate the second taxa is present). Since information is present on all taxa at any location with any data, there is little need to borrow strength across taxa (unlike the need to borrow strength across space to fill in missing areas and smooth over noise caused by limited data in each grid cell). This means that any downstream use of the results should avoid making use of the posterior covariance as an estimate of the correlation across taxa in the uncertainty of the estimates. Also note that one might scale the taxon-level point estimates to add to the total estimates, but there is no clear way to do this at the level of the posterior draws since the draws are computed independently between the total and all taxon fits.

In the GAM fits, we noticed some anomalies in the quasi-posterior draws for the occupancy model that were likely caused by numerical issues. In particular, for some taxa, there were draws of the occupancy probability that were more than five times as large as the probability point estimate. Most of these occurred for very low occupancy probabilities in areas outside the apparent range boundary for the taxa. There were also cases where draws of total

(non-taxon-specific) occupancy probability were near zero even though the probability point estimate was essentially one. As ad hoc, but seemingly effective solutions, we made the following adjustments to the draws:

1. set draws where the taxon-specific occupancy probability is greater than five times the point estimate to be equal to the point estimate, and
2. set all draws where the total point estimate was greater than 0.999 to be equal to 1.

#### Choice of smoothing and scale of averaging

We used cross-validation at the grid scale to:

1. choose between estimating potential biomass on the log-scale or original scale, and
2. determine the maximum number of spline basis functions, denoted as *k*.

With regard to the maximum number of basis functions, while the generalized additive modeling methods of Wood (2017) choose the amount of smoothing based on the data, using a large number of basis functions can result in slow computation. We chose to limit the number of basis functions (and thereby impose an upper limit on the effective degrees of freedom of the spatial smoothing estimated from the data), with that number informed by cross-validation. However, the imposed upper limit to the number of possible basis functions was large enough to have little effect on the amount of smoothing, although possibly imposing slightly more smoothing than without the limitation.

We used 10-fold cross-validation, randomly dividing the grid cells into 10 sets and holding out each set in turn. This allows us to assess the ability of the model to estimate biomass for cells with no data (and also gives us a good sense of performance for cells with very few points). Note that even with our incomplete sampling in Indiana and Illinois (Fig. 1), most unsampled grid cells are near to other grid cells with data. We considered values of ‘k’ in the set 100, 250, 500, 1000, 1500, 2000, 2500, 3000, and 3500.

The metrics used in cross-validation were squared error loss for the point predictions relative to the grid-level raw data and statistical coverage of prediction intervals of the grid-level raw data.

We calculated squared error weighted by the number of PLS points in the held-out cell and truncated both held-out values and predictions to maximum values of 600 Mg/ha to avoid having very large values overly influence the assessment. This also allows us to work on the original (not log) scale in our evaluation, as we don’t want to accentuate small differences at low biomass values.

We calculated 90% prediction interval coverage using a modified version of the quasi-Bayesian uncertainty procedure described previously. The modification to the sampling procedure involves drawing a random binomial value based on each draw of the occupancy probability, multiplied in turn by a random normal draw (exponentiated when fitting the model on the log scale) centered on the draw of the potential surface with variance equal to the residual variance from the potential model. The addition of the binomial draw and the residual variance produces a prediction interval for the data rather than the unknown process and allows us to assess coverage relative to an observed quantity. We calculated 90% prediction intervals using the 5th and 95th percentiles of the 250 draws for each held-out cell. Coverage was determined as the proportion of cells for which the observation fell into the interval, considering only grid cells with at least 60 PLS points. We also calculated the median length of intervals (and median log-length) to assess the sharpness of the intervals, as high coverage can always be trivially obtained from overly-wide intervals.

Unfortunately the coverage results cannot directly assess the uncertainty estimates provided in the data product, which provides intervals for gridded biomass. This is because the true biomass is unknown and thus cannot be used to judge coverage. We can only judge coverage of prediction intervals for the data. Thus, under- or over-estimation of uncertainty for the true quantities may be masked by compensating over- and under-estimation of the residual error of the data around the truth.

Cross-validation was done for total biomass and density as well as on a per-taxon basis.

## Results

### Model selection using cross-validation

The cross-validated weighted absolute error values for the biomass estimates can be seen in Table 2 for the model on the original scale and Table 3 on the log scale.

**Table 2:** Weighted absolute error for total biomass by number of basis functions for occupancy model (rows) and potential biomass model (columns) with potential model fit on original scale. Cells are weighted based on the number of PLS points in the cell.

|  | k for potential model |  |  |  |  |  |  |  |  |
| --- | --- | --- | --- | --- | --- | --- | --- | --- | --- |
| k for occupancy model | 100 | 250 | 500 | 1000 | 1500 | 2000 | 2500 | 3000 | 3500 |
| 100 | 38.47 | 36.35 | 34.91 | 33.81 | 33.43 | 33.34 | 32.58 | 32.54 | 32.51 |
| 250 | 38.05 | 36.02 | 34.60 | 33.51 | 33.14 | 33.05 | 32.30 | 32.26 | 32.23 |
| 500 | 37.75 | 35.67 | 34.29 | 33.23 | 32.87 | 32.78 | 32.04 | 31.99 | 31.97 |
| 1000 | 37.46 | 35.37 | 33.99 | 32.97 | 32.62 | 32.52 | 31.80 | 31.76 | 31.74 |
| 1500 | 37.35 | 35.24 | 33.86 | 32.85 | 32.49 | 32.40 | 31.67 | 31.63 | 31.61 |
| 2000 | 37.35 | 35.24 | 33.86 | 32.84 | 32.48 | 32.39 | 31.67 | 31.62 | 31.60 |
| 2500 | 37.17 | 35.07 | 33.68 | 32.66 | 32.31 | 32.22 | 31.50 | 31.45 | 31.43 |
| 3000 | 37.15 | 35.05 | 33.65 | 32.62 | 32.27 | 32.18 | 31.45 | 31.41 | 31.39 |
| 3500 | 37.15 | 35.05 | 33.65 | 32.62 | 32.27 | 32.17 | 31.45 | 31.41 | 31.39 |

**Table 3:** Weighted absolute error for total biomass by number of basis functions for occupancy model (rows) and potential biomass model (columns) with potential model fit on log scale. Cells are weighted based on the number of PLS points in the cell.

|  | k for potential model |  |  |  |  |  |  |  |  |
| --- | --- | --- | --- | --- | --- | --- | --- | --- | --- |
| k for occupancy model | 100 | 250 | 500 | 1000 | 1500 | 2000 | 2500 | 3000 | 3500 |
| 100 | 37.45 | 35.27 | 33.41 | 32.00 | 31.41 | 31.22 | 30.43 | 30.32 | 30.25 |
| 250 | 37.10 | 34.98 | 33.19 | 31.81 | 31.23 | 31.05 | 30.26 | 30.16 | 30.09 |
| 500 | 36.78 | 34.65 | 32.90 | 31.57 | 31.01 | 30.83 | 30.05 | 29.95 | 29.88 |
| 1000 | 36.52 | 34.38 | 32.65 | 31.38 | 30.83 | 30.65 | 29.88 | 29.78 | 29.71 |
| 1500 | 36.40 | 34.26 | 32.53 | 31.26 | 30.70 | 30.53 | 29.77 | 29.67 | 29.61 |
| 2000 | 36.39 | 34.26 | 32.50 | 31.23 | 30.68 | 30.51 | 29.75 | 29.65 | 29.58 |
| 2500 | 36.26 | 34.10 | 32.36 | 31.10 | 30.56 | 30.39 | 29.64 | 29.54 | 29.48 |
| 3000 | 36.24 | 34.08 | 32.33 | 31.08 | 30.53 | 30.36 | 29.62 | 29.52 | 29.46 |
| 3500 | 36.23 | 34.08 | 32.33 | 31.07 | 30.52 | 30.35 | 29.60 | 29.51 | 29.44 |

With regard to coverage, the models that fit the outcome on the original scale without log transformation for the potential model had a poor tradeoff of coverage and interval lengths. For k=2500 for occupancy and k=3500 for potential, for a 90% uncertainty interval, the coverage was 94.8% with a median interval length of 188, compared to coverage of 85.5% with a median interval length of 97 for the potential model on the log scale. While the 85.5% coverage is less than the desired coverage of 90%, we judge that the modest undercoverage is acceptable in light of the much shorter interval lengths. In addition, the uncertainty was roughly constant regardless of the value of the point estimate when working on the original scale, while the models using the log scale had uncertainty that increased with the size of the point estimate. This scaling of variance with mean (similar to that in a Poisson distribution) when using the log scale makes intuitive sense given the lower bound of zero.

Cross-validation results for total stem density are qualitatively similar to those for biomass with regard to how values vary with the number of basis functions (not shown). With regard to comparing results on the original and log scales, for k=2500 for occupancy and k=3500 for potential, the median absolute error was 41.5 and 40.1 for the original and log scales, respectively. Coverage was 91.5% and 85.4%, respectively, and the median interval lengths were 205 and 185, respectively.

Cross-validation results for taxon-level estimates are harder to interpret because there is one value per taxon. Also for grid cells outside the range limit of a taxon, estimates and intervals are generally very close to zero. As a result it is difficult to know how best to aggregate across taxa for summarization. The variation in cross-validation results with respect to the number of basis functions is qualitatively similar to the results for total biomass (not shown). For k=2500 for occupancy and k=3500 for potential, the average coverage (across taxa) of 90% uncertainty intervals was 97.8% for the original scale and 93.6% for the log scale, with a mean (across taxa) of median interval lengths (across cells) of 11.1 and 3.9 for the original and log scales, respectively.

Based on the cross-validation results we chose to fit models on the log scale. We also chose k=2500 for the occupancy models (for biomass, basal area, and stem density, and for total and taxon-level fitting) and k=3500 for the potential models. While values of k>2500 for occupancy reduced the estimated absolute error loss (i.e., improved the fits) slightly (see Table 3), larger k values increased computational time, so we chose to use k=2500. We did not assess k>3500. Based on the diminishing reductions in the loss as k increases beyond 2000 or 2500, it is unlikely that larger values of k would produce substantively important improvements in prediction.

### Estimated biomass and stem density

In Fig. 2, we show our estimated aboveground biomass and stem density, compared against the raw grid-level data (averaging over the point-level estimates), and with statistical uncertainty.

**Fig. 2.**
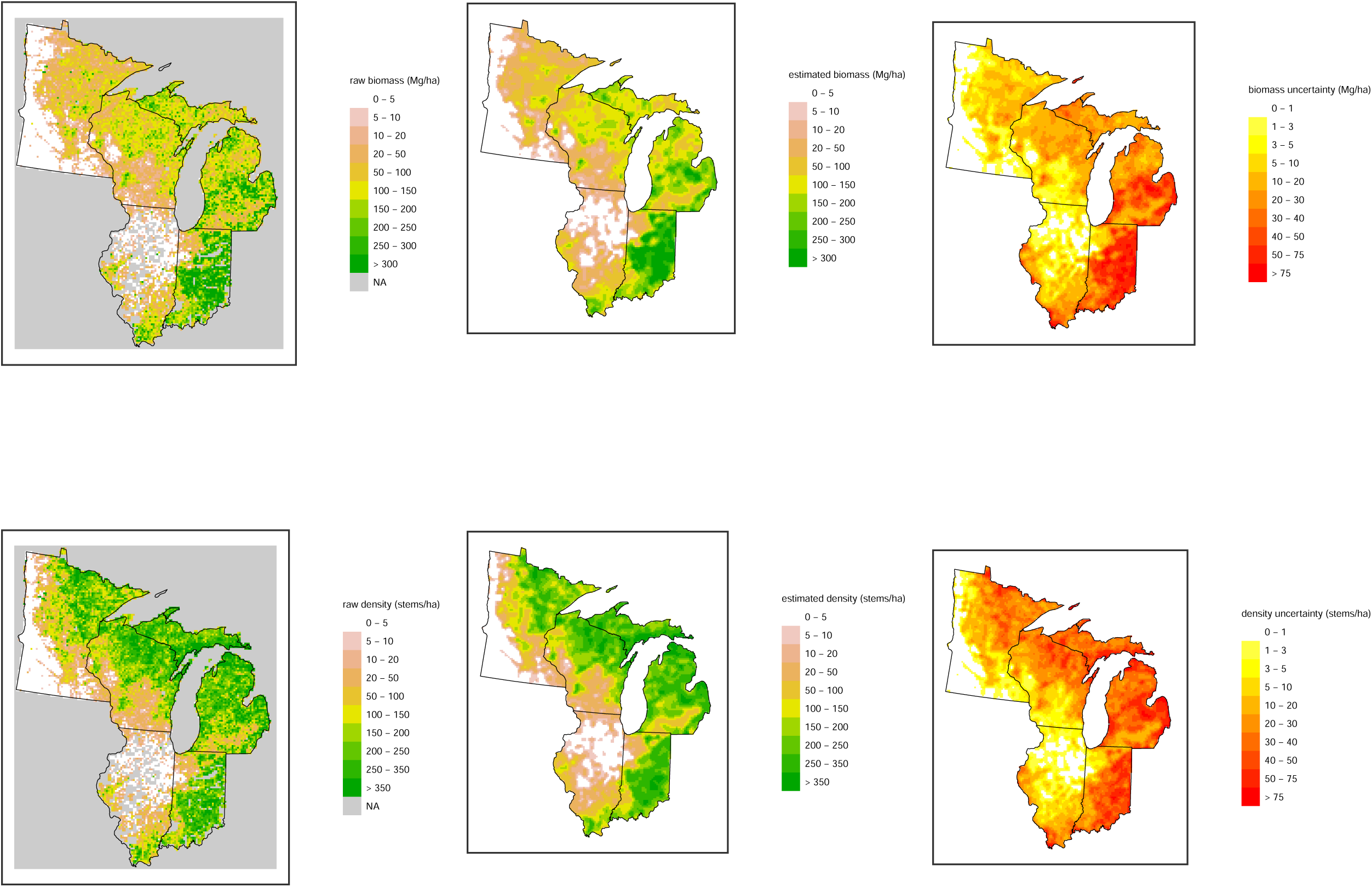
Raw data, predictions and uncertainty for total biomass (top row) and total stem density (bottom row). Point estimates from raw data in each cell based on the average point-level biomass (left column), predictions are estimates from the statistical smoothing model (middle column) and uncertainty estimates from the standard deviation of the quasi-Bayesian posterior draws (right column). Note that in the raw data plots, grey indicates data were not available for a grid cell (this occurs rarely, except in Illinois and Indiana).

Fig. 3 breaks the biomass down by a select set of taxa. Fig. 4 shows estimated biomass for all 21 taxonomic groups.

**Fig. 3.**
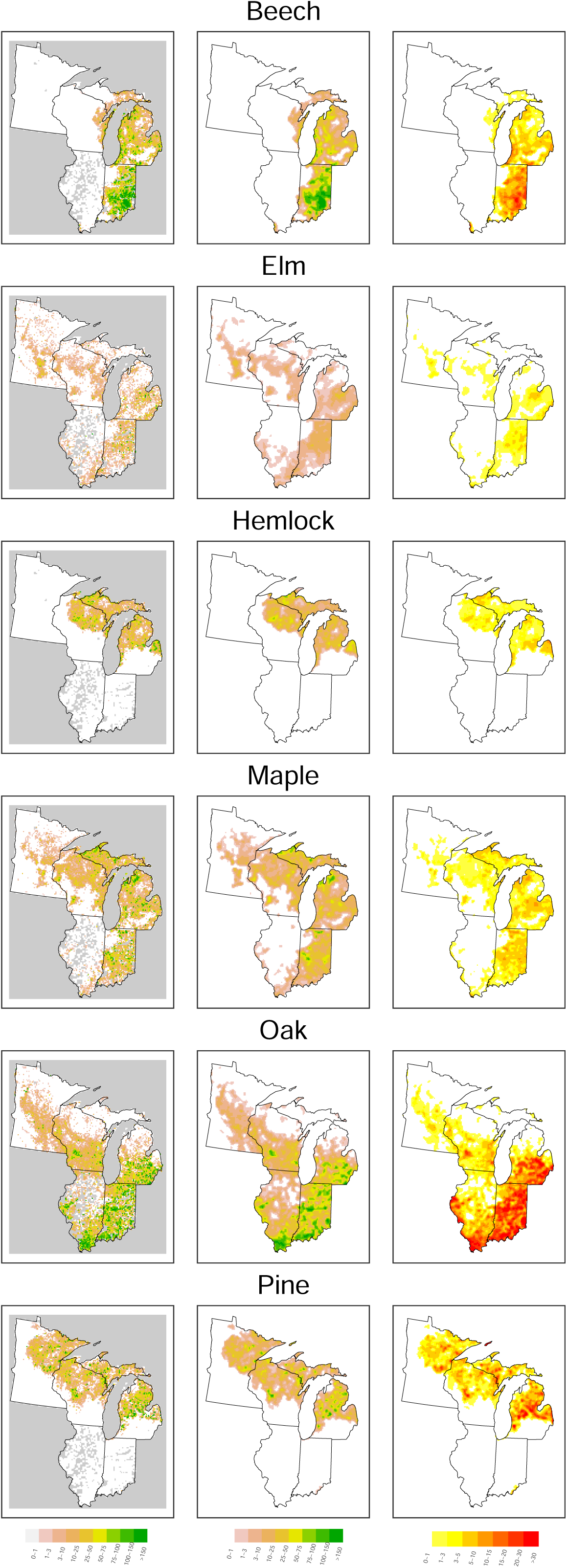
Raw data, predictions, and uncertainty for select taxa. Column ordering and figure formatting follow Fig. 2.

**Fig. 4.**
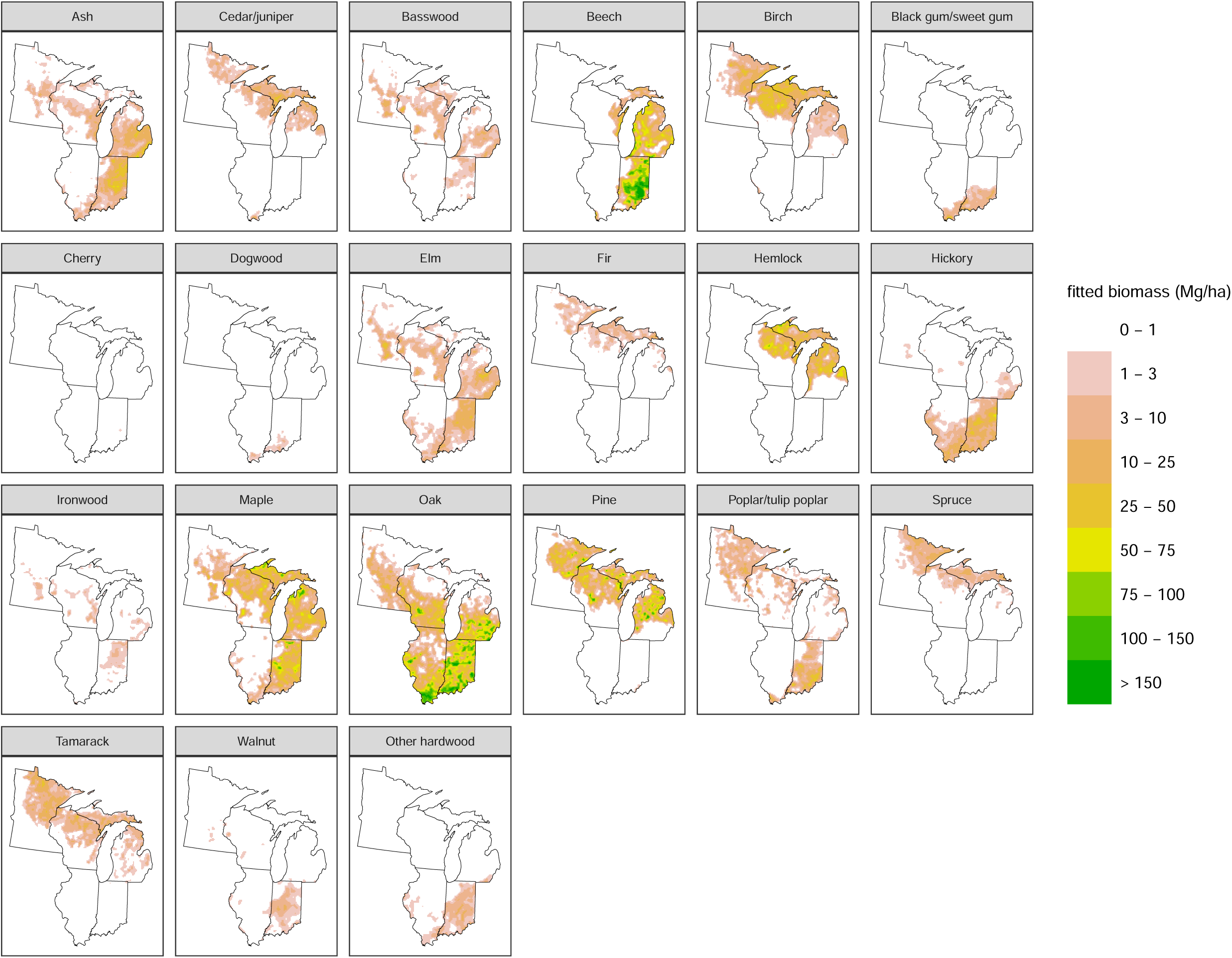
Predictions (point estimates) of biomass (Mg/ha) for all taxonomic groupings.

## Data product

We provide the following data products via the LTER Network Data Portal.

- Raw gridded stem density, aboveground biomass and basal area at https://portal.lternet.edu/nis/mapbrowse?scope=msb-paleon&identifier=26&revision=1 (DOI: 10.6073/pasta/801601af769fa5acade1ef07f6892bdd).
- Gridded statistically-smoothed stem density at https://portal.lternet.edu/nis/mapbrowse?scope=msb-paleon&identifier=24&revision=0 (DOI: 10.6073/pasta/1b2632d48fc79b370740a7c20a70b4b0).
- Gridded statistically-smoothed aboveground biomass at https://portal.lternet.edu/nis/mapbrowse?scope=msb-paleon&identifier=23&revision=0 (DOI: 10.6073/pasta/b246e05afb25dbe06b3006c5d18a4a2b).
- Gridded statistically-smoothed basal area at https://portal.lternet.edu/nis/mapbrowse?scope=msb-paleon&identifier=25&revision=0 (DOI: 10.6073/pasta/c3ae2363e4ae2e0f42a7c02b6f12b50a).

We will also soon provide point-level raw data for Indiana, Illinois, Michigan, and Minnesota. Point-level raw data for Wisconsin can be obtained by contacting co-author David Mladenoff.

The project Github repository (https://github.com/PalEON-Project/PLS_products) provides code for processing the point-level data and producing the data products above in the subdirectory named ‘R’. In the subdirectory ‘data/conversions’, we provide:

- our translation tables for translating surveyor taxon abbreviations to modern common names, including aggregation for the raw gridded values and statistical modeling done in this work,
- correction factors for the subregions of the domain for estimating point level tree density, and
- our assignments of allometric relationships for the PalEON taxa, based on Chojnacky et al. (2014) (also provided in Table 1).

## Discussion

We have presented high-resolution estimates, with uncertainty, of biomass, basal area, and stem density at the time of Euro-American settlement for a large area of the midwestern United States. These estimates can be used to answer ecological questions, as inputs for other analyses, and as a baseline for understanding changes in ecosystems, including carbon storage, under anthropogenic change.

While our estimates have a variety of strengths, including relatively high resolution, relatively uniform data density, coverage of a large area, careful data cleaning, and the use of statistical methods tailored to the data, there are of course limitations. The 8-km grid resolution prevents one from understanding variation at finer scales, including the stand level and variations from smaller scale effects such as local topography, including the effects of small fire breaks. For example, our total biomass and stem density estimates show a portion of the Minnesota River valley in southwestern Minnesota (see Fig. 2), but cannot resolve riparian forest (relative to grassland or upland forests) in smaller valleys. Our estimates smooth over the local variation, which can include sharp ecotone boundaries. In future work in this and other domains, we plan to make use of the point level data without initial gridding to try to estimate finer-scale variation, though one will always be limited by the natural resolution of the PLS survey points.

Our statistical model cannot represent range boundaries as it models variation in abundance as a continuously-valued spatial field with strictly positive (but often negligibly above zero) predicted biomass, basal area, and stem density, compounded by the smoothing mentioned above. Of course except in cases of distinct boundaries in environmental drivers that cause distinct range boundaries, range boundaries are generally fuzzy.

Our statistical model fits each taxon separately, for computational convenience and to limit the complexity of the spatial statistical models. Thus the uncertainty estimates do not capture any correlated uncertainty across taxa and analyses that aggregate estimates across more than one taxon (such as comparing two taxa or summing across multiple taxa) will not be able to correctly characterize uncertainty. For sums, one could, as we have done for total biomass, basal area, and density, sum the raw values and then apply the spatial statistical model. Finally, the sum across taxa of the taxon-specific estimates for a grid cell do not add to the estimate of total biomass, basal area, or stem density for that grid cell.

In this work, as in Paciorek et al. (2016) we chose not to use environmental covariates, such as soils, firebreaks, and topography (Grimm 1984, Shea et al. 2014), when estimating biomass, basal area, and stem density. Instead we limited our model to capture variation solely based on smoothing the data using Gaussian process techniques that rely on spatial distances. This avoids dependence on the environmental drivers of pre-settlement forest composition that might cause circular reasoning in subsequent analyses that use our data products. In addition, use of covariates could also lead to prediction that a taxa is present well beyond its range boundary in places where data are sparse.

The estimates and raw data are available as public data products, and our methods are fully documented with code available in our Github repository.

## Author contributions

CVC and CJP developed the statistical procedures for point-level estimation and spatial smoothing, respectively. JAP and CVC led the data cleaning and processing for Michigan, Illinois, and Indiana, while DJM provided the cleaned WI, MN and initial northern Michigan data. SJG and JWW led the initial processing of southern Michigan. JAP, CVC, and CJP did subsequent cleaning of data from both northern and southern Michigan. CJP and JAP wrote the code for data cleaning, data gridding and statistical estimation, with initial contributions from SJG and code/workflow review by AD. AD co-developed the workflow for biomass estimation using allometric equations. CJP wrote the paper with contributions from CVC and JAP and feedback from JWW, SJG, JSM, DJM, and AD.

## Acknowledgments

The authors are deeply indebted to all of the researchers over the years who have preserved, collected, and digitized survey records, in particular Jim Almendinger (Minnesota Department of Natural Resources), Robert McIntosh (formerly at University of Notre Dame), Ed Schools (Michigan State University Extension - Michigan Natural Features Inventory), and Ted Sickley (formerly at University of Wisconsin). We thank University of Wisconsin (Madison) undergraduates Madeline Ruid, Benjamin Seliger, Morgan Ripp and Daniel Handel for processing of the southern Michigan data and the Map Library in the Department of Geography at the University of Wisconsin for digitization of the Mylar maps. Indiana and Illinois data were made possible through the hard work of over 30 Notre Dame undergraduates in the McLachlan lab. We thank Jun Zhu, Xiaoping Feng, and Wesley Brooks for early work on the spatial statistical methods presented here. This work was carried out by the PalEON Project with support from the National Science Foundation MacroSystems Program through grants EF-1065702, EF-1065656, DEB-1241874 and DEB-1241868 and from the Notre Dame Environmental Change Initiative.

## Appendix A: Data collection and cleaning

### Wisconsin

Copies of the original field notes from Wisconsin are archived at the Wisconsin Board of Commissioners of Public Lands in Madison, Wisconsin. These records have been microfilmed and made available online (http://digicoll.library.wisc.edu/SurveyNotes/SurveyInfo.html or https://glorecords.blm.gov/default.aspx). The Wisconsin point data were digitized by the Mladenoff lab group, and have undergone several revisions over the last two decades in an effort to improve data accuracy (Radeloff et al. 2000, Manies and Mladenoff 2000, Mladenoff et al. 2002, Schulte et al. 2002, Liu et al. 2011,). This constitutes the cleaned raw data used in this work, which can be obtained by contacting David Mladenoff.

We then processed the cleaned raw data as follows.

We excluded 4088 points with the first tree marked as ‘QQ’ by the Mladenoff lab as this indicates the presence of water at the point. We used the vegetation code for each point (documented at the Wisconsin DNR website) to exclude points that might induce a negative bias in our estimates because trees were missing because of standing water. Specifically, we excluded 2803 points with one tree that are marked as Creek, Marsh, Swamp, Lake, and River. Note that we included points marked as Wet Prairie or “low land, low wet area”, judging these areas to be terrestrial, albeit often wet.

Three areas southwest of Green Bay had no data because they were Menominee Native American lands and were not surveyed (see Fig. 1). We excluded these points and few other points in Wisconsin (a total of 736 points) for which there was no information on trees at the point. Later surveys on these lands are not included here.

There were 670 points that were missing survey years. The survey year for these was imputed based on nearby points.

### Minnesota

Copies of the original submitted field notes from Minnesota are at the Minnesota Historical Society in St. Paul, Minnesota. These records have been digitized by three projects (Grimm 1981, Almendinger 1985, Minnesota County Biological Survey 1996 [unpublished]) and are available online at the Minnesota DNR Bearing Tree Database (https://gisdata.mn.gov/dataset/biota-original-pls-bearing-trees). More details are given in Almendinger (1997). The data used in this work were obtained from the Minnesota DNR by the Mladenoff lab group in earlier work. This constitutes the cleaned raw data used in this work and provided in the data product.

We then processed the cleaned raw data as follows.

We used the vegetation code for each point (documented in Almendinger (1997)) to exclude points that might induce a downward bias in our estimates because trees were missing because of standing water. Specifically, we excluded 20560 points with no trees or one tree that are marked as Creek, Marsh, Swamp, Lake, and River. We retained 24243 points with two to four trees in such areas as it is hard to know how much of the area surrounding these corners is under water; this may lead to some downward bias in our density estimates in Minnesota. We excluded 902 points with missing taxon information and missing ecotype as these appeared unusable. Most occurred in far northern Minnesota (the Boundary Waters area) or along straight east-west lines, suggesting problems with the points. We excluded 1560 Forest, Grove, Bottom and Pine grove points with missing taxa information for all trees, as this is inconsistent with the presence of forest. We included points marked as Wet Prairie, judging these to be terrestrial, albeit often wet.

### Northern Michigan

Copies of the original submitted field notes from northern Michigan are at the Michigan State Archives in Lansing, Michigan. These records have been microfilmed and are available online (http://seekingmichigan.org/discover/glo-survey-notes or https://glorecords.blm.gov/default.aspx). Michigan surveyor observations for the Upper Peninsula of Michigan and the northern section of the Lower Peninsula (see Fig. 1 of Goring et al. (2016)) were digitized by the Mladenoff lab group in earlier work. Co-authors CVC and JP added additional points in Ontonagon, Schoolcraft and Gogebic Counties. The northern Michigan data were further processed to keep one record for locations that had two georeferenced data entries with identical tree information. In cases where there were two georeferenced data entries in the same location, with either a) one entry providing tree information and the other no tree information or b) both entries having the same tree information except for one attribute (typically the bearing), the point with no information or with less information, respectively, was removed.

The exact point coordinates for Isle Royale appeared to be incorrect (some points are in Lake Superior) and some points appeared to be duplicated. We omitted all data from Isle Royale in the current analysis, but in future work we plan to re-enter the data from the original survey notes.

From initial spot checks in Dickinson County we determined tree diameter and distances were transposed in the data transcription in many cases; these were corrected. Spot checks also indicated that distances in Iosco County (and scattered points in other counties) that had been listed with decimal values needed to be converted from chains to links (multiplied by 100) to standardize with the rest of the database. There were some trees with outlying diameter values greater than 48 inches dbh. All of these were checked carefully in the original field notes and were corrected if necessary.

The cleaned raw data represent the data obtained and processed as described just above and provided in the data product. We then processed the cleaned raw data as follows.

Unlike in Minnesota and Wisconsin, we do not have vegetation codes, so we cannot distinguish one-tree points that occur because of water from one-tree points in areas with low density. Some points have qualitative notes but in general these do not indicate the presence of any water. All one-tree points were retained, but they may contribute to a downward bias in density and biomass.

There were 2810 points with no trees indicated (i.e., with no taxa noted) for which we attempted to determine if these were truly points that had been surveyed that had no nearby trees. All such points (1316 points) without any surveyor notes were excluded. We included points where the notes indicated no trees (e.g., ‘no witness trees’, ‘no trees convenient’, ‘no other tree data’) but excluded points where the notes indicated water (102 points), lost information (73 points) or that the tree was used as the corner or no other trees were recorded (874 points). In the latter case, we cannot compute a density estimate (it would be infinity) for a point with one tree at zero distance.

### Southern Michigan

Copies of the original submitted field notes from southern Michigan are at the Michigan State Archives in Lansing, Michigan. These records have been microfilmed and are available online (http://seekingmichigan.org/discover/glo-survey-notes or https://glorecords.blm.gov/default.aspx). Field notes were transcribed to topographic maps allowing the points to be displayed geographically and were then converted to Mylar maps by Denis Albert and Patrick Comer of the Michigan Natural Features Program for Albert et al. (2008). Ed Schools provided these Mylar maps temporarily to the Williams lab, who digitized them to a point-based ArcGIS shapefile and co-authors CVC and JP conducted a number of checks described below. When necessary they used the original field notes to check and make corrections.

There were some townships with no survey points (visible as small areas with fewer points per cell, albeit not zero points because the township and grid cell borders are not aligned, in south-central and southeastern Michigan in Fig. 1). Spot checks indicate these are generally caused by missing data from the original surveys, with the original field notes unavailable for unknown reasons.

We removed points already contained within the southern Michigan dataset. An initial assessment of the data digitized from the mylar maps indicated that the diameter and distances were transposed during digitization; we corrected these. When the Mylar maps were created, points on the township boundaries were entered twice for approximately 4000 exterior township corners (mainly on the southern and eastern township borders), resulting in four trees noted per corner when only two were surveyed. We kept the two of the four trees listed when they were located in quadrants inside the township. We excluded 469 points in cases where there was ambiguity because the trees were in quadrants outside the township. We plan to obtain data from these points from the original PLS field notes in future processing. An additional 68 interior township survey points had 3-4 trees but for which only two trees were truly surveyed. These were checked against the original PLS field notes and corrected. For entries with azimuths less than zero or greater than 360 we checked the survey notes and corrected as needed. There were some trees with outlying diameter values of greater than 48 inches dbh. All of these were checked carefully in the Mylar maps and/or original field notes and were either corrected or retained only if they were clearly noted in either source.

The cleaned raw data represent the data obtained and processed as described just above and provided in the data product. We then processed the cleaned raw data as follows.

Due to extensive incomplete data on the Mylar maps, 27 townships in the Detroit region (primarily Monroe and Lenawee Counties, with one township in Washtenaw) were re-entered and replaced by the McLachlan lab using the same protocol as used for the Indiana and Illinois data.

The Mylar maps for many areas of southern Michigan (outside of the Detroit region) have no quarter-section points. The areas with quarter-section points tend to be in savanna / low-density areas. Given this selection bias, we removed all 6593 quarter-section points that were present. This can be seen in the marked decrease in points per cell in the southern portion of the lower peninsula of Michigan in Fig. 1.

We excluded 40 points with three trees because surveyors in this area were instructed to only marked two trees. These points may be those with a corner tree plus two additional trees but extracting valid data from these points would require looking at the field notes.

Unlike in Minnesota and Wisconsin, we do not have vegetation codes, so we cannot distinguish one-tree points that occur because of water from one-tree points in areas with low density. Some points have qualitative notes but in general these do not indicate the presence of any water. All one-tree points were retained, but they may contribute to a downward bias in density and biomass.

### Indiana and Illinois

Copies of the original submitted field notes from Indiana and Illinois are at the Indiana State Archives in Indianapolis and the Illinois State Archives in Springfield. These records have been microfilmed and are available online in the National Archives (https://catalog.archives.gov/id/566714). Data from Indiana and Illinois were purchased from the Indiana State Archives (Commission on Public Records, Indiana State Archives, Indianapolis) Indiana) and Hubtack Document Resources (http://hubtack.com/ab/index.php), respectively, and processed by the McLachlan lab. Data entry for these states is ongoing. Originally, townships to digitize were chosen to provide an even distribution across both states. Since then, specific areas of each state (e.g., the Kankakee watershed, the Yellow River watershed, the savanna-closed forest transition north to south in Illinois and west to east in Indiana, and locations of US Forest Service Forest Inventory plots) have been chosen to complement ongoing projects in the lab. PLS land notes are transcribed by undergraduates in the lab. These data are then subjected to an initial QA/QC check by the original readers, followed up by a second QA/QC check by a different individual in the lab, georeferenced to the section and quarter-section locations and finally, reviewed for a final set of QA/QC checks. The R code used for the QA/QC checks and georeferencing are available in Github at: https://github.com/PalEON-Project/IN_ILTownshipChecker.

The cleaned raw data represent the data obtained and processed as described just above and provided in the data product. We then processed the cleaned raw data as follows.

There were 18 points in the Illinois data near the Illinois-Wisconsin border that were very close to points in Wisconsin. These were removed to avoid near-duplication of points. A small number of points were missing survey years. The survey year for these was imputed based on nearby points.

Points recorded as Water (where surveyor nodes indicated standing water) or having no data (which is not the same as having no trees) were omitted. Points recorded as Wet (where surveyor notes indicated water was ephemeral) were included as these were judged to be terrestrial. All one-tree survey points are retained as the survey information (including qualitative notes by the surveyors) indicates that such points were because of low density and not the presence of water.

### Other notes

We excluded 127 points in Minnesota, Wisconsin, and northern Michigan in which at least one tree was marked as dead (or ‘dry’) or where the taxon was judged ‘indeterminable’ by the Mladenoff lab. In almost all these cases, there would be fewer than two live trees after excluding the dead or unknown cases.

## Appendix B: Correcting the Morisita plotless density estimator

Deputy surveyors were instructed (ca. 1805) to mark “two or more adjacent trees in opposite direction as nearly as may be” at each section or quarter-section corner (White 1983). The procedure was implemented from the earliest PLS surveys in Illinois as evident by two witness trees invariably being recorded. Later the survey instructions required that witness trees be in different sections; this two-tree procedure remained unchanged on exterior and interior lines through the 1850s in Ohio, Illinois, Indiana, and Michigan. In 1846 the instructions for Wisconsin and Iowa (including Minnesota) changed to include four witness trees at township and section corners and two trees in opposite sections at quarter-section corners (White 1983).

The predominant sampling design in the Midwest PLS was two-tree corners (84.8%). Starting in 1786, exterior corners in Ohio, southern Illinois before 1810, all of Michigan, southern Wisconsin before 1845, Indiana after 1829, and Illinois after 1838 were sampled with two trees on same side inside the township (2sH). In northern Wisconsin after 1846 and all of Minnesota this 2sH design was applied at only exterior section corners. At the remaining exterior and all interior corners in all Midwestern states the sampling design generally followed trees in opposite halves or dominant “opposite” with a mixture from “adjacent” quadrants (2oH or 2S). Four-tree corners (9% of all Midwest) were prevalent only from northernmost Illinois (16%) after 1837, northern Wisconsin (14%) after 1846, and all of Minnesota (24%) after 1847. Virtually all these four-tree samples were at section corners with 38% on the township’s exterior lines. The majority (61%) of the four-tree corners of the upper Midwest were at interior section corners and 95% had one witness tree in each quadrant/section.

In summary, there were three common sampling designs in the Midwest: two trees on the inside of exterior lines (2sH), two trees in somewhat opposite halves on interior lines (2oH or 2S), and four trees at section corners at township corners and on both exterior and interior lines.

Given this heterogeneity, we corrected the Morisita estimator to minimize error due to different sampling geometries and several known surveyor biases (Manies et al. 2001, Kronenfeld and Wang 2007, Bouldin 2008, Liu et al. 2011, Williams and Baker 2011, Hanberry et al. 2011, Hanberry et al. 2012a, Hanberry et al. 2012b), as discussed in Goring et al. (2016). Our approach allows for spatial variation in surveyor methods by applying correction factors based on the empirical sample geometry, and known surveyor biases deviating from this design (Cogbill et al., in progress). These estimates are based on empirical examination of the underlying data, and have been validated using simulations on stem-mapped stands (Cogbill et al., in progress). The correction factors vary by state (and in some cases regions within a state), date, internal versus external point, section versus quarter-section, and two-versus four-tree points and can be found in our Github repository. There are four correction factors: kappa accounts for the sampling design (two-versus four-tree points and where the two trees are located relative to quadrants and halves), theta for sector bias, zeta for azimuthal censoring, and phi for inclusion of trees less than 8 inches dbh, as discussed in Goring et al. (2016) and Cogbill et al. (in progress).

